# Dissecting the co-segregation probability from genome architecture mapping

**DOI:** 10.1101/2022.08.15.503981

**Authors:** Lei Liu, Xinmeng Cao, Bokai Zhang, Changbong Hyeon

## Abstract

The genome architecture mapping (GAM) is a recently developed methodology that offers the co-segregation probability of two genomic segments from an ensemble of thinly sliced nuclear profiles, enabling to probe and decipher the 3D chromatin organization. The co-segregation probability from GAM, which typically probes the length scale associated with the genomic separation greater than 1 MB, is, however, not identical to the contact probability obtained in Hi-C, and its correlation with inter-locus distance measured with FISH is not so good as the contact probability. In this study, by using a polymer-based model of chromatins, we derive a theoretical expression of the co-segregation probability as well as that of the contact probability, and carry out quantitative analyses of how they differ from each other. The results from our study, validated with in-silico GAM analysis on 3D genome structures from FISH, suggest that to attain strong correlation with the inter-locus distance, a properly normalized version of co-segregation probability needs to be calculated based on a large number of nuclear slices (*n* > 10^3^).

**SIGNIFICANCE:** By leveraging a polymer model of chromatin, we critically assess the utility of co-segregation probability captured from GAM analysis. Our polymer model, which offers analytical expressions for the co-segregation probability as well as for the contact probability and inter-locus distance, enables quantitative comparison between the data from GAM, Hi-C, and FISH. Although the plain co-segregation probabilities from GAM are not well correlated with inter-locus distances measured from FISH, properly normalized versions of the probability calculated from a large number of nuclear profiles can still reasonably represent the inter-locus distance. Our study offers instructions of how to take full advantage of GAM analysis in deciphering 3D genome organization.

## INTRODUCTION

Among a number of experimental methods to decipher the three-dimensional (3D) chromosome/genome structure at high resolution (1–13), a recently developed genome architecture mapping (GAM) (8, 9, 14), enabling genome-wide mapping of chromatin contacts, has gained much attention. In GAM, cells are first fixed and cryosectioned, and next processed through laser microdissection to produce ultrathin nuclear slices, called nuclear profiles (NPs). The sequencing of the DNA content in each NP allows one to identify genomic loci present in the NPs and calculate their frequencies (14) (Fig. 1A). Loci that are distant in space are expected to have smaller chance to be co-sectioned in the same NP. It has been presumed that the co-sectioning/co-segregation probability between two genomic sites (*c*_*ij*_), *i* and *j*, is inversely related to their mean spatial distance 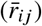 (14).

**Figure 1:**
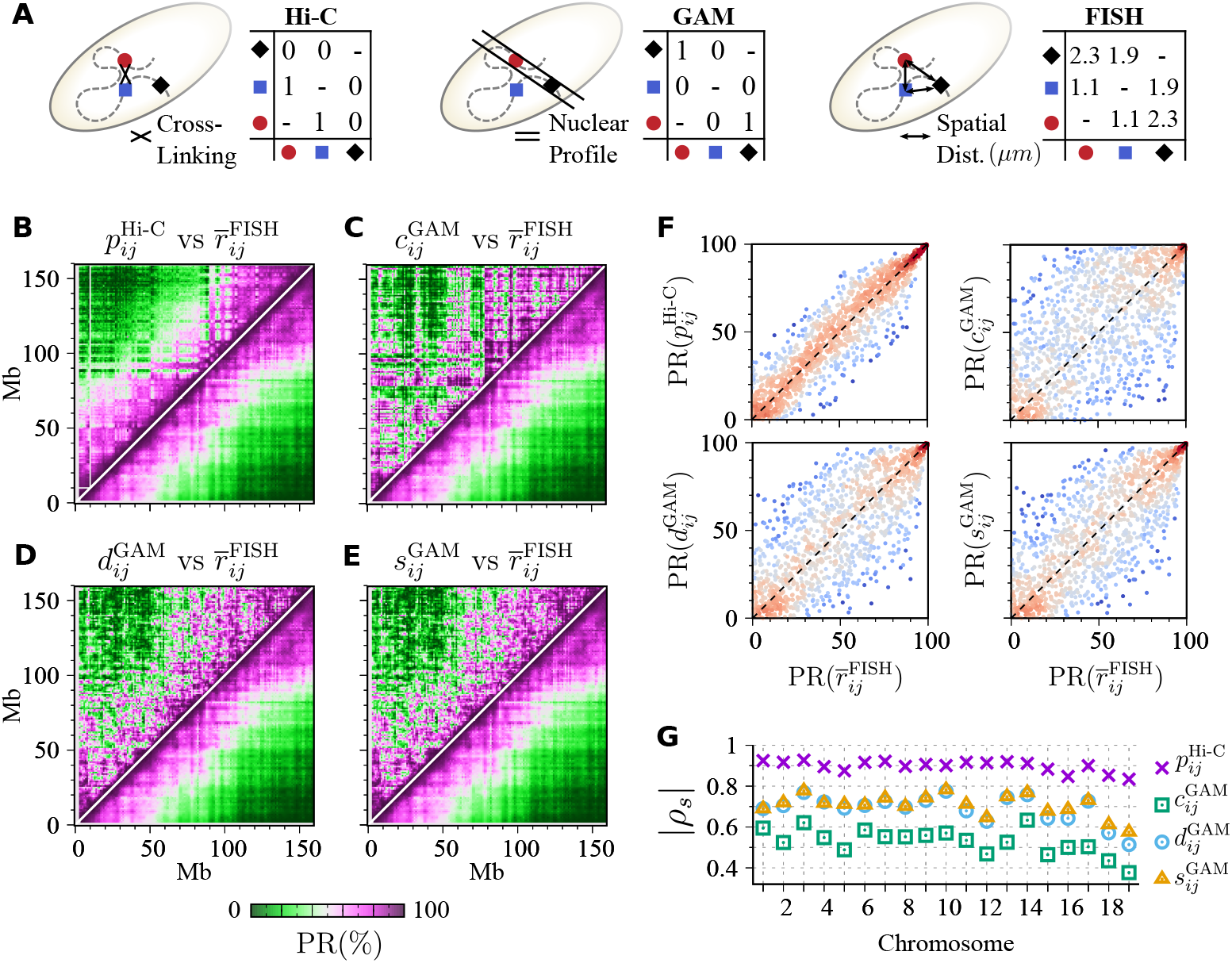
Comparison between GAM, Hi-C, and FISH. **(A)** Schematics illustrating the three methods. From an ensemble of cells, Hi-C measures the cross-linking frequencies between two genomic loci; GAM measures the co-segregation frequencies in a nuclear profile; FISH measures the inter-locus spatial distances. **(B-D)** GAM data are compared with FISH that probed the chromosomes of mouse ESC using DNA seqFISH+ method (24), which offers the 3D coordinates of 2,460 loci spaced approximately 1 Mb apart across the whole genome in 446 cells. Heatmaps of the percentile rank (PR) of the mean inter-locus distance 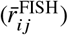 on the chromosome 3 binned at 1 Mb (right, bottom corner of each panel) (24) versus **(B)** PR of Hi-C contact probability (5) 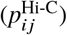, **(C)** PR of co-segregation probability from GAM (14) 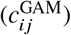, **(D)** PR of normalized linkage disequilibrium (14) 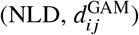), and **(E)** PR of normalized point-wise mutual information (27) 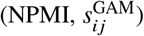, each of which is demonstrated on the top left corner of the panel. **(F)** The scatter plots of PRs calculated in (B)-(E). The density of data point is color-coded from blue to red. **(G)** The Spearman’s rank correlation coefficient (|*ρ*_*s*_ | = −*ρ*_*s*_) of 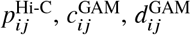 and 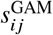 against 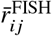 for the 19 autosomes of mouse ESCs.

GAM has several key advantages over C-based techniques in that it is ligation-free (12, 15). Furthermore, in contrast to Hi-C which requires millions of sample cells, only hundreds of cells may suffice for GAM to generate a robust genome-wide co-segregation map. Given that clinical samples are often limited in number and given in a sectioned form, GAM can be more practical than other methods when studying disease-related genome reorganization. While not easily accessible in Hi-C, a number of important properties of 3D genome, such as higher-order chromatin contact (16–21), lamin and nuclear body association (22–26), can be measured by leveraging cryoFISH-combined GAM.

The potential of GAM is built upon the premise that the data acquired from the mapping faithfully reflect the 3D organization of genome. To demonstrate the utility and fidelity of GAM, Beagrie *et al*. (14) used co-segregation frequencies (or probabilities, *c*_*ij*_) to identify topologically associated domains (TADs) that are interacting with each other. Referencing to cryoFISH images, they showed that the interacting TADs have higher contact probabilities (*p*_*ij*_) and smaller spatial distances 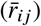 than non-interacting TADs. More recently, by using the Strings and Binders Switch model of chromatin, Fiorillo *et al*. (27) reported almost perfect correlations between *c*_*ij*_ and 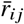 with a Spearman’s ranking correlation coefficient of ρ_*s*_ < −0.98 in four regions of genomic sizes < 7 Mb. However, given that those results were obtained with a limited number of samples, how statistically general is the inverse relationship between 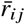 and *c*_*ij*_ still remains as an open question. To be specific, when the co-segregation probability of a particular chromatin segment pair is greater than another pair (*c*_*ij*_ > *c*_*kl*_), can we assert that their mean distances always satisfy the inequality 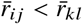?

To address the above question, we calculate 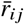 by using 3D imaging data of mouse embryonic stem cell (ESC) (24), and compare them with *c*_*ij*_ obtained from GAM (24) by means of their percentile rank (PR), i.e., PR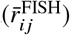 vs. PR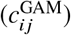, where the superscripts “FISH”, “GAM” and “Hi-C” are added to specify the experimental data source for clarity. For the case of 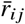, we consider that PR is higher when 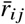 is smaller. As shown in Fig. 1B, PR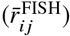 depicted in a matrix form for the chromosome 3 displays a pattern similar to the PR of the contact probability *p*_*ij*_ from Hi-C. However, for the cases of GAM-based co-segregation probability (*c*_*ij*_) and its variants, – the normalized linkage disequilibrium (NLD, *d*_*ij*_) and the normalized point-wise mutual information (NPMI, *s*_*ij*_) (27) (see Materials and Methods) – such similarity between the patterns is significantly weaker (Fig. 1C-E). The scatter plot of PR(*p*_*ij*_) against PR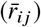 also displays stronger correlation than those of PR(*c*_*ij*_), PR(*d*_*ij*_), and PR(*s*_*ij*_) (Fig. 1F). The Spearman’s rank correlation of *p*_*ij*_ with 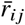 is significantly greater than that of GAM-based measures (*c*_*ij*_, *d*_*ij*_, *s*_*ij*_) for all autosomes of mouse ESCs (Fig. 1G).

In the light of the general agreement between Hi-C and FISH (3, 24, 28–31), the significantly weaker correlation of co-segregation frequencies from GAM with the mean inter-locus distances from FISH, demonstrated in Fig. 1, is rather surprising. To better understand the meaning of data from GAM, we consider an analytically tractable, simple polymer model, representing chromosomes inside a nucleus, and derive co-segregation probability (*c*_*ij*_), which shows that the *c*_*ij*_ changes with the coverage, the thickness, and the number of nuclear slices. We validate our theoretical predictions from the polymer model by carrying out in-silico GAM analysis on a publicly available 3D genome structure dataset. Our study, which enables quantitative comparison between GAM, Hi-C, FISH measurements, offers instructions of how to take full advantage of GAM analysis in deciphering 3D genome organization.

## MATERIALS AND METHODS

### GAM, Hi-C, FISH and DamID datasets

The genome-wide GAM data was downloaded from GEO repository (14) (GSE64881). It contains binary information, denoting either the absence or the presence of chromatin segments binned at 1 Mb, in each of 408 NPs, for all autosomes of mouse ESCs. For each chromosome, we counted the frequency of the *i*-th segment and the frequency of the (*i, j*) segment pair in the same profile, which yields the segregation and co-segregation probability, *c*_*i*_ and *c*_*ij*_, respectively.

We used the Hi-C dataset of mouse ESC measured by Dixon *et al*. (5). KR-normalized intra-chromosome contact probability at 1Mb resolution, *p*_*ij*_, was calculated by using the Juicer toolbox (32).

The DNA seqFISH+ dataset was fetched from Zenodo database (https://zenodo.org/record/3735329), where 3D coordinates of genetic loci spaced approximately 1 Mb apart across the whole genome in 446 cells were deposited. The genomic coordinates of the imaged loci were converted from mm10 to our reference genome assembly mm9 with the UCSC Genome Browser utility liftOver (33). Following Takei *et al*. (24), we separated two homologous chromosomes in each mouse ESC based on the consensus of the spectral and hierarchical clustering of imaged loci. For each intra-chromosome loci pair (*i, j*) binned at 1 Mb, we then calculated their spatial distances for each allele in all cells, which yields the mean inter-locus distance 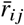.

The DamID of Lamin B1 protein in mouse ESCs was downloaded from GSE17051 (34), where the base-2 logarithm of the fold enrichment of the interactions between chromatin loci and nuclear lamin (*q*_LB1_) was available. A chromatin segment has a positive (negative) value of *q*_LB1_ if it has higher (lower) chance to associate with lamin than the genome-wide average level.

### Two variants of co-segregation probability

Of several possible methods for normalizing GAM data (35), we consider the two most popular ones, normalized linkage disequilibrium (NLD) and normalized pointwise mutual information (NPMI).

i. To account for the observation that different loci have different chances to be cryosectioned, the normalized linkage disequilibrium (NLD), which was originally proposed in population genetics to calculate the non-random association of two alleles at different loci (36, 37), has been employed for the analysis of GAM (14). The NLD is defined as

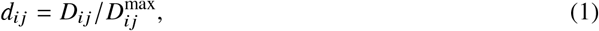

where

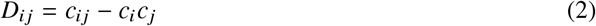

compares the co-segregation probability of *i* and *j*-th loci (*c*_*ij*_) to the probability of statistical independence, normalized by the theoretical maximum

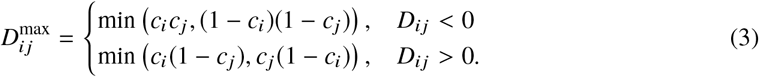
ii. Alternatively the normalized pointwise mutual information (NPMI) has been used for the GAM analysis as well (27):

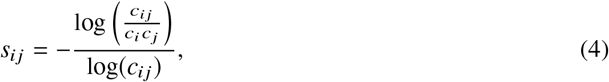

where the distance of cosegregation probability of *i* and *j*-th loci is measured in logarithmic scale (log *c*_*ij*_) from that of the statistical independence (log *c*_*i*_*c*_*j*_) and normalized by log *c*_*ij*_.

The two quantities, *d*_*ij*_ and *s*_*ij*_, bounded between −1 and 1, are conceptually similar in that both measure the distance of the joint probability of cosegregation from the case of statistical independence (i.e., *c*_*ij*_ = *c*_*i*_*c*_*j*_); however, they differ from each other in that the measurement by the *d*_*ij*_, which amounts to the covariance (or correlation), is restricted to linear relationships, while *s*_*ij*_ can capture more general relationship between two random variables (38).

## RESULTS AND DISCUSSIONS

### Theoretical analyses

#### Heterogeneous loop model

We use the Heterogeneous Loop Model (21, 39–45) (HLM), to derive analytical expressions for the contact and co-segregation probabilities as well as mean inter-locus distances, and study the relations (or correlations) between them. In HLM, chromatin fibers are modeled as a linear polymer chain composed of *N* coarse-grained segments, each with a prescribed genomic size. We assume that the effective energy potential of chromatin can be described by a sum of harmonic restraints on the spatial distances between all segment pairs,

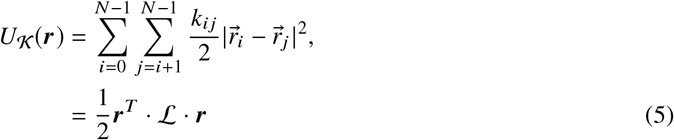

where 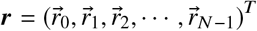 specifies the 3D structure of the polymer chain, and the *N* × *N* Laplacian matrix L is defined as ℒ = 𝒟 − 𝒦 where 𝒦 is a stiffness matrix of element *k*_*ij*_ and 𝒟 represents a diagonal matrix with 𝒟_*ii*_ = ∑_*j*_ *k*_*ij*_. The probability of the chromatin to adopt a particular structure is then written as

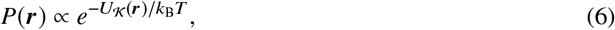

where *k*_B_*T* is the energy unit of the model with *k*_*B*_ the Boltzmann constant and *T* the absolute temperature. The model parameters (𝒦) for a genomic region of interest can be determined based on Hi-C data.

The greatest advantage of HLM is that structural and dynamic properties of chromosome can be directly derived based on Eqs. S1 and 6 along with Hi-C data. HLM and its variants have been exploited to study experimental measurements (21, 40–45). The contact probability calculated from the HLM is in excellent agreement with measurements (21, 41, 42). Specifically, despite the cell-to-cell variability of 3D genome over population (45, 46), the intra-chromosomal inter-locus spatial distance distributions predicted by HLM can still be validated against those measured from DNA seqFISH+ imaging (see Fig. 1 in Ref. (21)).

Although it does not affect the discussion in this work, the HLM has limitations when it gets to a resolution greater than 𝒪(10^2^) base pairs, where each monomer represents the scale of 1-2 nucleosomes, whose interactions (bending, torsion, stacking, etc.) can no longer be effectively approximated by harmonic restraints.

#### Contact probability and spatial distance between two genomic segments

After transforming 𝒦 into a covariance matrix Σ whose matrix element is denoted as *σ*_*ij*_ ≡ (Σ)_*ij*_, one finds the mean spatial distance between the *i*- and *j*-th segments averaged over all possible chromosome configurations as (detailed derivations are given in the Supplementary Information)

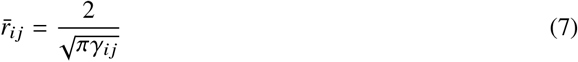

With 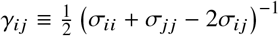, and their contact probability

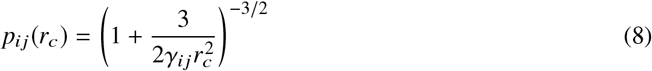

where *r*_*c*_ is the effective capture radius of the cross-linking agent, and is the only tunable parameter of the contact probability in Hi-C.

In this theoretical framework, the two observables (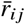 and *p*_*ij*_) are related as 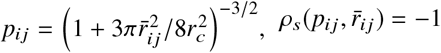, and hence they are in perfect correlation, giving rise to the Spearman’s rank correlation coefficient 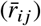. The consistency between Hi-C (*p*_*ij*_) and FISH 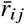 shown in Fig. 1B and G has been demonstrated by a number of experimental studies (28–30, 47).

In addition, we trained HLM for each autosome of mouse ESCs by using their Hi-C data binned at 1 Mb, and calculated contact probability and mean inter-locus distance based on Eq. S5 and S4, respectively. As shown in Fig. S1, not only *p*_*ij*_ from HLM is highly correlated with that from Hi-C (*ρ*_*s*_ ;:: 0.95), but also 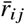 from FISH can be well predicted with the correlation coefficient *ρ*_*s*_ ≃ 0.9, which is even slightly higher than the correlation between Hi-C and FISH data. These results suggest that HLM is a proper 3D model of chromosome.

#### Co-segregation probability

The probability of the *i*-th genomic segment being in a horizontal slice 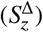 with a thickness Δ sectioned at a height *z* relative to the center of mass (COM) of the chain is derived as follows (see Supplementary Information)

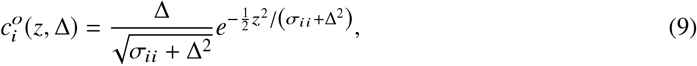

and the co-segregation probability is formulated as

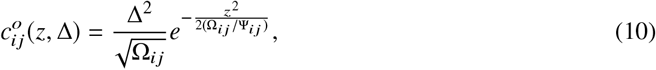

where

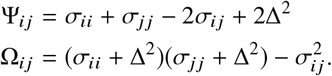

Eqs. S4-S11 are the general theoretical expressions derived in the framework of HLM with *σ*_*ij*_ (or the force parameters *k*_*ij*_) left unspecified. Their validity can be examined against a Gaussian phantom chain consisting of 20 monomers, which is a special case of HLM where the condition of *k*_*ij*_ = 1 for |*i* − *j*| = 1 and *k*_*ij*_ = 0 otherwise is assigned. By numerically generating an ensemble of 100,000 polymer chain configurations whose COM is restrained to the origin of the coordinate system (Fig. 2; see Supplementary Information for numeric details), we counted the frequency of the *i*-th monomer 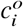 (or the joint frequency of the *i*- and *j*-th monomers, 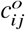) in a horizontal slice 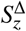. The numerical results of (co-)segregation probability obtained from explicit 3D structures (Fig. 2A) are in perfect agreement with our analytic expressions in Eqs. S10 and S11 (Figs. 2B and C).

**Figure 2:**
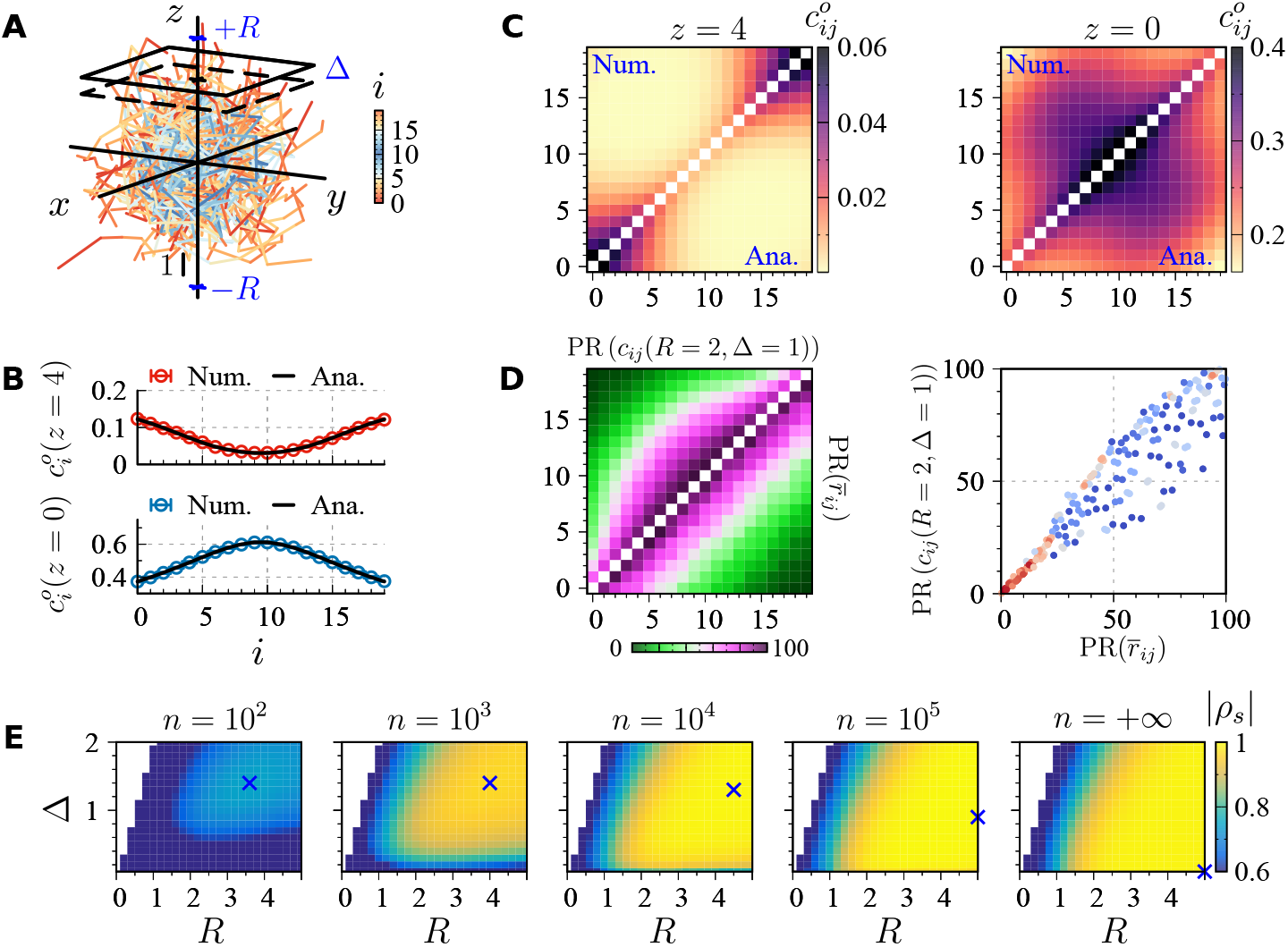
Co-segregation of monomers in a Gaussian phantom chain. **(A)** Depicted are 100 chains randomly selected from an ensemble, each composed of 20 monomers. GAM data would be collected from an ensemble of slices with a thickness Δ, and at a height *z* (|*z*| < *R*). The terminal and the central parts of the chain are colored in red and blue, respectively. **(B)** Segregation and **(C)** co-segregation probabilities, 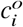 and 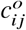, calculated at different values of *z* with Δ = 1. **(D)** (Left panel) Percentile rank (PR) of the co-segregation probability (top left) compared with the PR of the mean inter-locus distance (bottom right). (Right panel) Scatter plot of (PR *c*_*ij*_), PR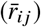). The density of data point in each pixel is color-coded from blue to red. **(E)** Spearman’s rank correlation coefficient (*ρ*_*s*_) between *c*_*ij*_ and 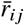 as a function of *R* and Δ, where the co-segregation probabilities were calculated by using *n* slices. The |*ρ*_*s*_ | with *n* → ∞ is obtained by using Eqs.S4 and S14. The condition of (*R*^∗^, Δ^∗^) that gives rise to the strongest correlation |*ρ*_*s*_ | for different *n* is marked with the symbol ×.

Eqs. S10 and S11 make it explicit that the (co-)segregation probability not only depends on the thickness, but also on the location of the slice. As expected, 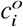 and 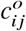 increases with the thickness of the slice. But, their variation with *z* is nontrivial. The radius of the polymer ensemble depicted in Fig. 2A is *R* ≈ 5. Thus, in a NP sectioned at *z* = 4, the chain segments located at the terminals have a higher odds to be sliced than those around the center, and this trend is reversed in another NP sectioned at *z* = 0 (Fig. 2B).

As shown in Fig. 2A, the chain segments colored based on their position along the chain contour (blue for the segments around the center and red for the segments at the two ends) visualize the origin of the *z*-dependencies. Due to the geometrical restraint on the COM of an individual chain, the chain terminals lie at the periphery, and hence the distribution of chain segments is radially *non-uniform*. The ranking order of 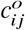 demonstrates qualitative difference with *z* as well (Fig. 2C).

In GAM, NPs are collected from many samples, i.e., random slices in position and orientation over many nuclei (cells). Provided that the slice range of nuclei is [−*R, R*], the mean segregation probability averaged over the uniform distribution of *z* ∈ [−*R*, +*R*], which corresponds to data resulting from the GAM analysis for a large number of slices (*n* ≫ 1), would be obtained as

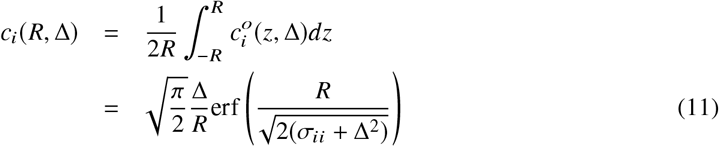

with 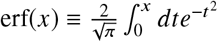. Similarly, the mean co-segregation probability is obtained as

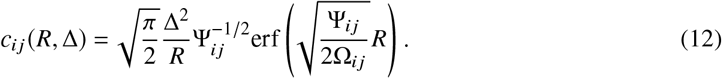

Eq. S14 suggests that *c*_*ij*_ is decided by the range of the nuclei being sliced, namely [−*R*, +*R*]. Unlike Ψ_*ij*_ which is a monotonic function of the mean distance, 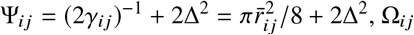 changes non-monotonically with 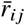. Thus, the presumption of a monotonic relation between *c*_*ij*_ and 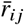 not hold in the outcomes derived from HLM. As illustrated in Fig. 2D, PR *c*_*ij*_ clearly differs from PR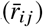.

To demonstrate the effect of the sample size (*n*) on the Spearman’s rank correlation between *c*_*ij*_ and 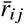, we divided the ensemble of polymer chain structures into 100, 000/*n* replicas, calculated *c*_*ij*_ for each replica, and plotted the replica-averaged value of |ρ_*s*_| as a function of *R* and Δ in Fig. 2E. As *n* increases from *n* = 10^3^ to *n* → ∞, the value (*R*^∗^, Δ^∗^) that maximizes |ρ_*s*_| shifts towards large *R* and small Δ.

The segregation (Eq. 11) and co-segregation probabilities (Eq. S14) were derived by assuming that the sectioning was made with Gaussian probability at height *z* and that the NPs were collected uniformly over the nuclear volume. They can be derived by assuming other models of the sectioning probability and slice position profile; however, the resulting probabilities remain qualitatively the same (see the text and Figs. S2 and S3 in the Supplementary Information).

#### Co-segregation probability of HLM-based chromosome model

The behaviors of co-segregation probability obtained from Gaussian phantom chain are sufficiently general, but can be made more realistic. HLM of 1-Mb genomic region on chromosome 5 of GM12878 cells, trained in reference to its Hi-C data (Fig. 3A), produces an ensemble of structures characterized with three distinct domains (Fig. 3B). As found from the Gaussian phantom chain model (Fig. 2), we confirm the dependence of the (co-)segregation probability on the position of the slice (i.e., the dependence of 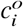 and 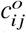 on *z*; see Figs. 3D, E), the nonuniform radial distribution of chromosome (Figs. 3C, D, E), and the improved correlation between *c*_*ij*_ and 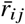 at greater *n* (Figs. 3F, G).

**Figure 3:**
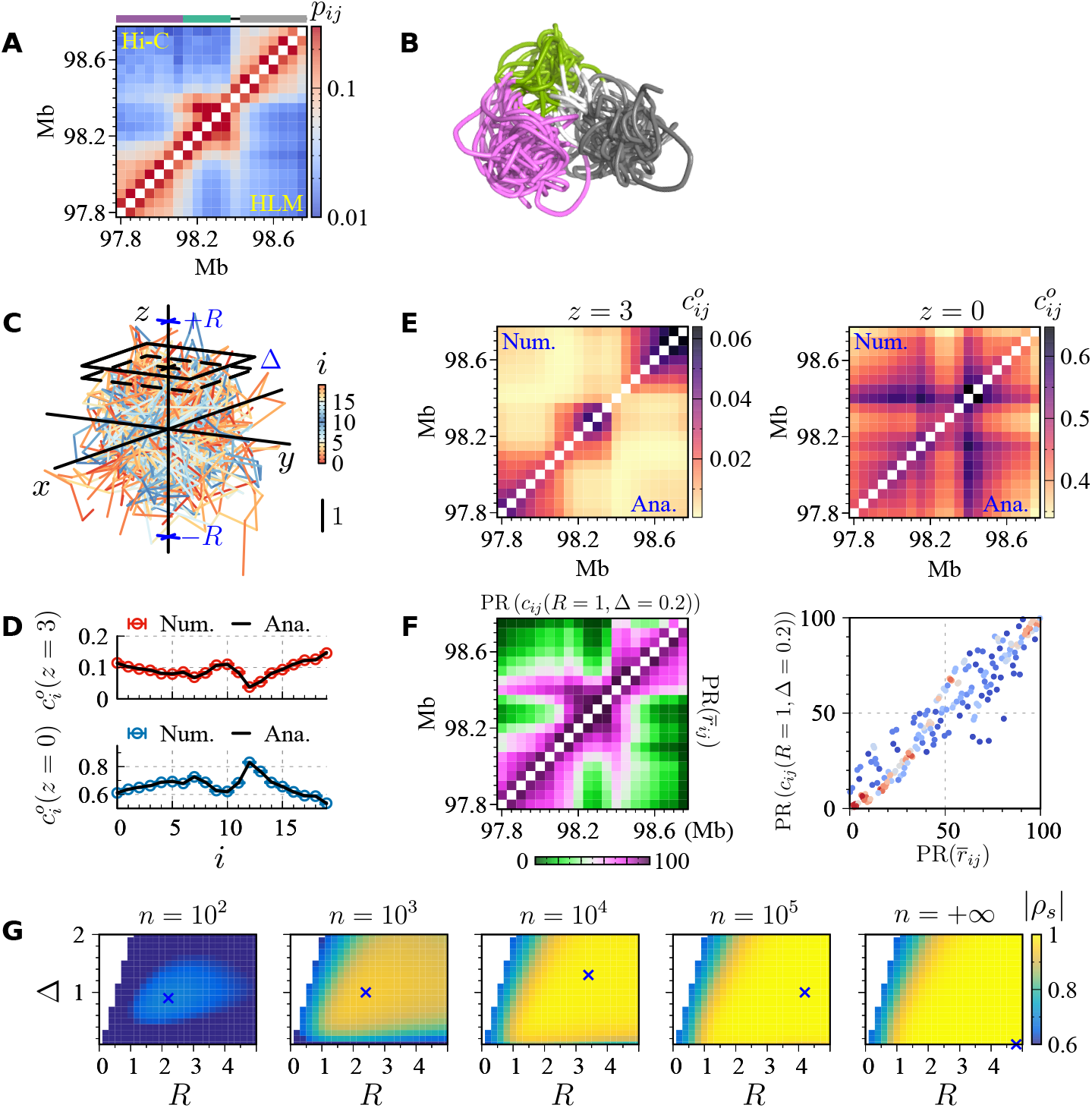
Co-segregation probability from the HLM-based structural ensemble at a resoluton of 50 kb for a 1-Mb genomic region on chromosome 5 (chr5:97,800,000-98,800,000) of GM12878 cells. **(A)** The contact probability (*p*_*ij*_) measured by Rao *et al*. using Hi-C (6) (top-left) is compared with that predicted by HLM (bottom-right). Details about the determination of model parameters based on Hi-C data and reconstruction of 3D chromosome structures can be found in the Refs. (21). **(B)** An ensemble of chromatin chains (*N* = 30) randomly selected from the most populated state of the structural ensemble. Color-coded are three distinct domains. **(C)** An ensemble of 100 randomly selected chromatin structures without alignment. Co-segregation was determined from an ensemble of slices of a thickness Δ, and a stochastic height *z* satisfying |*z*| < *R*. **(D)** Segregation and **(E)** co-segregation probabilities, 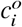 and 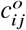, calculated at different values of *z* with Δ = 1. **(F)** (Left panel) PR of the co-segregation probability (top left) compared with that of the mean inter-locus distance (bottom right). (Right panel) Scatter plot of (PR (*c*_*ij*_), 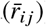). **(G)** Spearman’s rank correlation coefficient (*ρ*_*s*_) of *c*_*ij*_ against 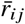 as a function of *R* and Δ. The symbol × marks the condition of (*R*^∗^, Δ^∗^) that gives rise to the strongest correlation at different *n*.

### Correlation of Hi-C and GAM against FISH

#### In silico GAM analysis of FISH data

The interactions between chromatin segments can be mediated by many nuclear compositions such as nucleoli, lamins, granules, and so forth. Although such contributions may implicitly be accounted in the energy potential of HLM (Eq. S1), efficacy of our theory for the 3D structure modeling of whole genome remains unexplored. To this end, we next performed in-silico GAM analysis on the 3D genome structures measured by Takei *et al* (24).

We randomly sampled 100,000 genome structures with replacement from the mouse ESC dataset of the DNA seqFISH+ experiment, and then translated the COM of each structure to the origin, and rotated the structure to a random orientation. Horizontal slices with a thickness Δ were obtained from the structures by randomly sampling the position along the *z*-axis in the range of *z* ∈ [−*R*, +*R*] (Fig. 4A). The resulting ensemble of slices were partitioned into replicas, each containing *n* slices.

**Figure 4:**
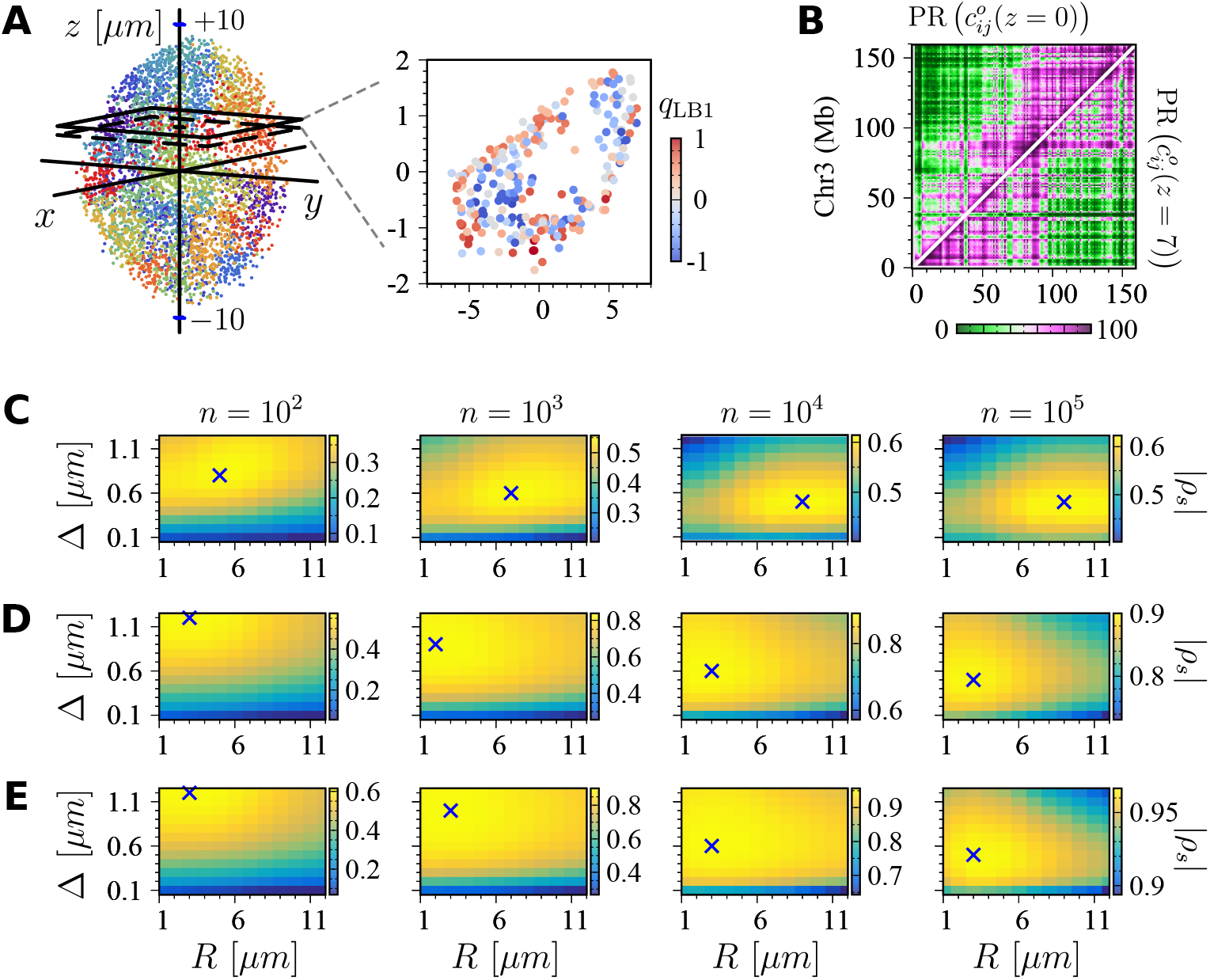
In silico GAM on the DNA seqFISH+ dataset (24). **(A)** A typical 3D image of the genome of mouse ESC where different chromosomes are distinguished by different colors. Shown on the right is a slice, in which the sectioned loci are color-coded by their lamin association propensities (*q*_LB1_). **(B)** PR of *c*_*ij*_ sectioned by slices with Δ = 0.5 *µ*m and at *z* = 0 and 7 *µ*m, shown in the top-left and bottom-right panel, respectively. The Spearman’s rank correlation coefficient between the two 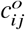’s is *ρ*_*s*_ = 0.83. The Spearman’s rank correlations (|*ρ*_*s*_ |) of 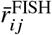 against 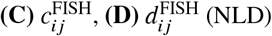, and **(E)** 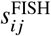 (NPMI) as a function of the slice range of nuclei (*R*) and slice width (Δ) with varying sample size (*n*), where the “×” symbol marks the value of (*R*, Δ) that maximizes |*ρ*_*s*_|. The heatmaps in **(C-E)** quantify the variations of correlation over the full range of their respective data.

First, as demonstrated by the different lamin colocalization propensities of loci shown in Fig. 4A, the chromatin segments inside the cell nucleus has a non-uniform radial distribution. Second, we confirm that co-segregation probabilities 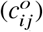 calculated at the different positions of the nuclear slice (*z* = 0 and 7 *µ*m) are qualitatively different from each other (Fig. 4B). More pronounced dependence of 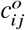 on *z* is found for other chromosomes (Figs. S4A and S5A). Third, the Spearman’s rank correlation of 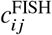 with 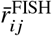 follows the same trend with *R*, Δ, and *n* (Fig. 4C) in accord with our theoretical analyses using the Gaussian polymer chain (Fig. 2E) and HLM for chromosome 5 of GM12878 cell (Fig. 3G). For *n* ≫ 1, the correlation of three co-segregation probabilities with 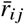 is maximized at a large *R* and at a small Δ. However, it is noteworthy that even with *n* = 100, 000, the best correlation between 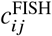 and 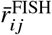 is about |*ρ*_*s*_| ≃ 0.63, which is significantly weaker than the correlation of *p*_*ij*_ of Hi-C with 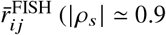 see Fig. 1G). The other two *c*_*ij*_-related quantities, NLD and NPMI, show better correlations with 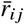 than *c*_*ij*_; however, the number of NPs should be greater than 10^3^ for the maximal values of |*ρ*_*s*_| for NLD and NPMI to exceed 0.9 (Fig. 4D and E). Similar conclusions are drawn from chromosomes 12 (Fig. S4) and 18 (Fig. S5).

#### Stratified comparison of GAM with FISH

Thus far, all intra-chromosome segment pairs were taken into account to calculate the *ρ*_*s*_. To see how the fidelity of GAM data in representing the chromosome architecture changes at different genomic scales, we re-calculated ρ_*s*_ against the mean spatial distance 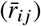 from FISH (24) by restricting our analysis to the loci pairs separated by a certain genomic distance (48, 49). Despite large variations among different chromosomes (Fig. 5A and Fig. S6), at short genomic separation (|*i* − *j*| < 40 Mb), Hi-C contact probability (*p*_*ij*_) is significantly better correlated with 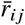 than the GAM-based co-segregation probability and its variants (*c*_*ij*_, *d*_*ij*_, *s*_*ij*_). Only for the extremely long-range segment pairs (|*i* − *j* | > 75 Mb), the GAM dataset marginally outperforms the *p*_*ij*_ from Hi-C. Given that 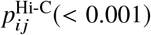 is typically orders of magnitude smaller than the corresponding 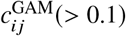 for long-range pairs, the bad performance of Hi-C might be an outcome of its relatively higher NSR (Eq. 14).

**Figure 5:**
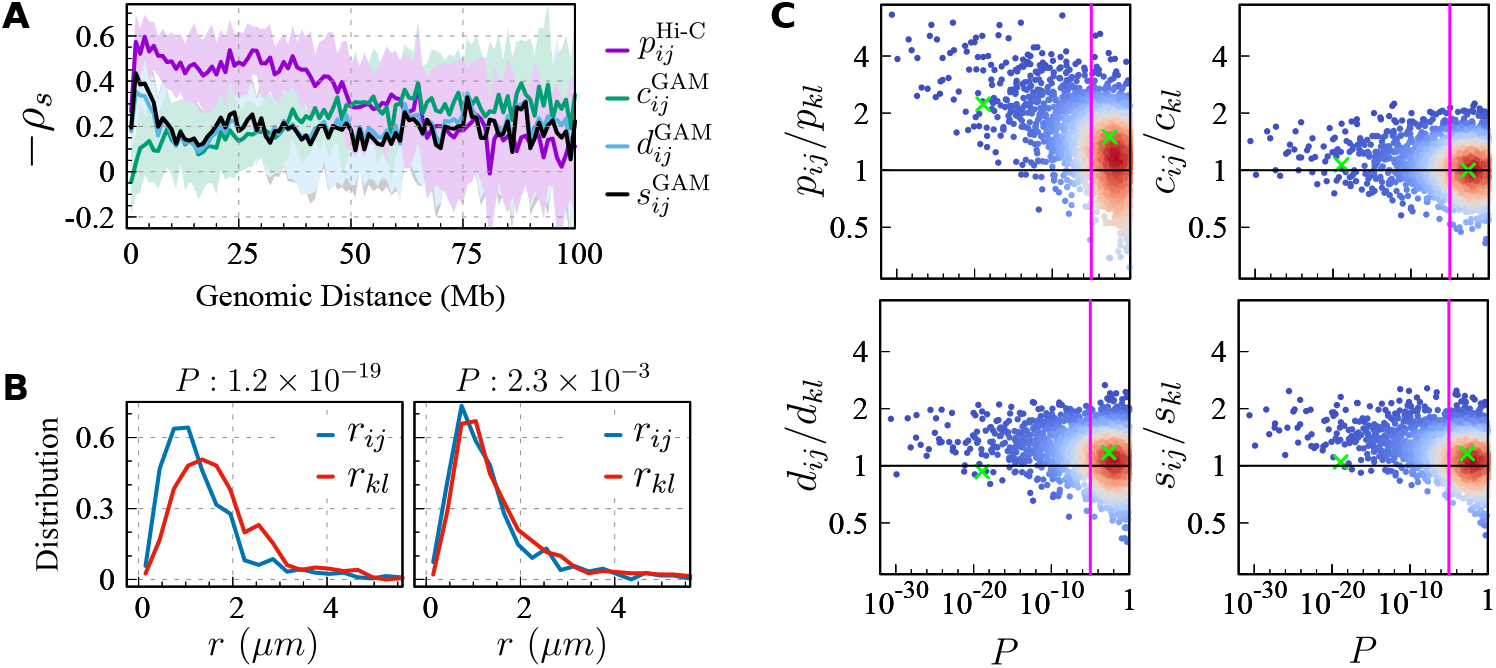
Stratified comparison of GAM (14) and Hi-C (5) against FISH data (24). **(A)** Autosome-averaged Spearman’s rank correlation coefficient (|ρ_*s*_|) between 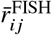 and Hi-C contact probability (*p*_*ij*_), GAM co-segregation probability (*c*_*ij*_) and its two variants (*d*_*ij*_, *s*_*ij*_) as a function of the genomic separation between the two sites (*i, j*). The mean and the standard deviation of *ρ*_*s*_ averaged over all autosomes are plotted with solid lines and shades, respectively. **(B)** Comparison between the distance distributions of one intra-chromosomal pair *i, j* (blue) and another (*k, l*) based on FISH. *P*-value to reject the hypothesis (*r*_*ij*_ < *r*_*kl*_) calculated from Mann-Whitney U test is shown on the top. The left (right) panel shows an example with a small (large) *P* value. **(C)** Four possible ratios of a segment pair (*i, j*) to another (*k, l*) versus their *P*-value. The two examples in (B) are marked with the symbol (×). The vertical line depicts *P* = 1.0 × 10^−5^.

Lastly, for two intra-chromosomal pairs on the chromosome 3 separated by 4 Mb, say (*i, j*) and (*k, l*), we calculate a *P*-value between the distributions of two inter-locus distances from FISH (24) to assess the statistical significance of the statement that the spatial distance of a particular pair is shorter than another (*r*_*ij*_ < *r*_*kl*_) instead of merely comparing their means (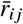 versus 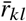). When the *P*-value is smaller, it is more likely that the two pairs display different distances (Fig. 5B); if *r*_*ij*_ < *r*_*kl*_, then *p*_*ij*_ > *p*_*kl*_ is highly likely. However, according to our explicit calculation summarized in Fig. 5C, the above statement regarding the relation between inter-locus distance and contact probability does not nicely translate into the GAM-based co-segregation probabilities. In Fig. 5C which plots the data points *x*_*ij*_/*x*_*kl*_, where *x*_*ij*_ denotes either the contact (*p*_*ij*_) from Hi-C or one of the GAM-related co-segregation probabilities (*c*_*ij*_, *d*_*ij*_, or *s*_*ij*_) as a function of their *P*-value, we find that the number of data points satisfying *x*_*ij*_/*x*_*kl*_ > 1 among those with *P* < *P*^∗^ = 1 × 10^−5^, i.e.,

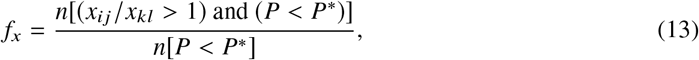

is *f*_*p*_ = 0.93, *f*_*c*_ = 0.65, *f*_*d*_ = 0.84, and *f*_*s*_ = 0.87, leading to *f*_*p*_ > *f*_*s*_ ≳ *f*_*d*_ ≫ *f*_*c*_. The *p*_*ij*_ from Hi-C reflects the inter-locus distance more faithfully than the GAM-related co-segregation probabilities. Although the plain co-segregation probability from GAM (*c*_*ij*_) is poorly correlated with *r*_*ij*_, the properly normalized versions of co-segregation probabilities, NLD (*d*_*ij*_) and NPMI (*s*_*ij*_), can still reasonably represent the inter-locus distances (*r*_*ij*_).

### CONCLUDING REMARKS

In practice, the sample size (*n*) in GAM is finite. With the mean (*c*_*ij*_) and variance 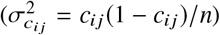 of the co-segregation probability, the corresponding noise-to-signal ratio (NSR) is given by

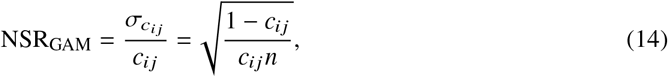

which decreases with *n* as 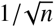 (27). It is of note that the NSR of contact probability (*p*_*ij*_) has a form identical to Eq. 14 when *c*_*ij*_ is replaced with *p*_*ij*_. For the contact and co-segregation probabilities, which display power-law decays (*p*_*ij*_, *c*_*ij*_ ∼ |*i* − *j*|^−*α*^) (50), measurements are less reliable (or large NSR) for long-range segment pairs (Eq. 14). Furthermore, since the variations of the co-segregation probability with respect to the variations of Δ and *R* satisfy ∂*c*_*ij*_/∂Δ < 0 and ∂*c*_*ij*_/∂ *R* > 0 for *n* ≫ 1 (Eq. S14), the NSR_GAM_ is reduced for *δ*Δ < 0 and *δR* > 0, i.e.,

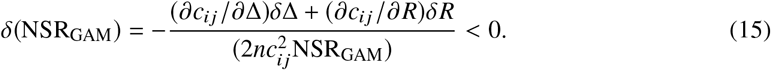

The reduced NSR_GAM_ at larger *R* and smaller Δ accounts for better correlation with 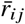, which manifests itself as the enhanced |*ρ*_*s*_| for *n* ≫ 1 under the same condition (Figs. 2E and 3G).

Meanwhile, HLM predicts that the inter-locus distance (*r*_*ij*_) measured by FISH has a mean and variance of 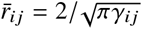 and 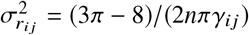, respectively. Thus, NSR of *r*_*ij*_ from FISH is

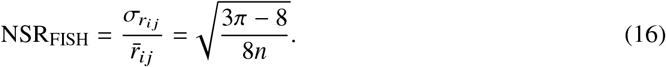

Comparison of Eq. 16 with Eq. 14 (NSR_FISH_ < NSR_GAM_) suggests that for a given sample size, the distance measurement using FISH is more precise and reliable than GAM (or Hi-C) when the co-segregation probability *c*_*ij*_ (or contact probability, *p*_*ij*_) is less than 8/3*π* ≈ 0.85.

To recapitulate, our theory predicts that unlike the contact probability, the co-segregation probability change non-monotonically with the spatial distance between two loci, which accounts for the discrepancy between GAM and FISH data on chromosome scales (Fig. 1). We demonstrate by using Gaussian phantom chain (Fig. 2), HLM of chromosome (Fig. 3), and in-silico analysis of experimental genome structures (Fig. 4) that the co-segregation probability depends on the slice range of nuclei, the thickness and the number of nuclear sections. The correlation between the co-segregation probability from GAM and the spatial distance between two genomic segments is not perfect, but it is only moderate both at chromosome-wide (Fig. 1) and at short genomic range (Fig. 5). As a result, GAM data is not always straightforward to interpret. Thus, the findings made with GAM analysis, such as inter-chromosome contact, higher-order interaction, and associations with lamin and nucleolus, are usually supplemented with other measurements (9, 14). Our theoretical analysis of GAM based on HLM shows that if (co)-segregation probabilities from NPs with slice thickness of ∼ 0.5 *µm* are only available data, then as long as the data are collected from a sufficient number (*n* ≳ 10^3^) of NPs NPMI (*s*_*ij*_), albeit not perfect, is slightly better suited than NLD (*d*_*ij*_) for faithful interpretation of the 3D chromatin organization.

## AUTHOR CONTRIBUTIONS

L.L. and C.H. designed the research. L.L. carried out all simulations. L.L., X.C., B.Z., and C.H. analyzed the data and wrote the article.

## ACKNOWLEDGMENTS

L.L. thanks Xue Sun for helpful discussions. This work was financially supported by Zhejiang Provincial Natural Science Foundation, National Natural Science Foundation of China, Zhejiang Sci-Tech University (LQ22B040001, 12104404, 20062226-Y to L.L.) and Korea Institute for Advanced Study (CG035003 to C.H.). We thank the Center for Advanced Computation in KIAS for providing computing resources.

## SUPPLEMENTAL INFORMATION

The expressions of mean inter-locus distance and the contact probability between chromatin segment pairs of Heterogeneous Loop Model (HLM) are derived in Sec. 1 and 2, respectively (39–43), followed by the detailed derivation of the co-segregation probability in Sec. 3. In the last section, we describe numerical generation of a 3D structural ensemble of the model.

### 1 MEAN INTER-LOCUS DISTANCE

The chromatin fiber in a genomic region of interest was modeled as a linear polymer chain composed of *N* coarse-grained monomers each representing a chromatin segment with a prescribed genomic length. The interaction energy of the fiber has a form of

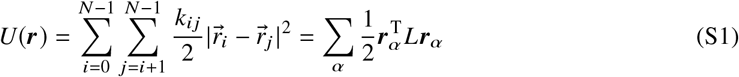

where **r**_*α*_ = (*r*_0,*α*_, *r*_1,*α*_, *r*_2,*α*_, · · ·, *r*_*N*−1,*α*_)^*T*^ and the subscript α represents *x, y* and *z*. The Laplacian matrix ℒ is defined as ℒ = 𝒟 − 𝒦, where 𝒦 is a stiffness matrix of elements *k*_*ij*_ and 𝒟 is a diagonal matrix of elements 𝒟_*ii*_ = ∑_*j*_ *k*_*ij*_.

Upon translating the center of mass (COM) of the polymer to the origin of the coordinate system, ℒ can be transformed to a new matrix,

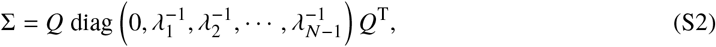

where the *i*-th column of the orthorgonal matrix *Q* and *λ*_*i*_ are the corresponding *i*-th eigenvector and eigenvalue of ℒ, respectively. By using the notation of

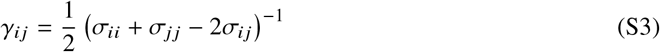

where *σ*_*ij*_ is the (*i, j*)-th element of Σ, the mean distance and the mean squared distance between the *i*- and *j*-th sites in 3D space, that can be measured by FISH imaging, are obtained as

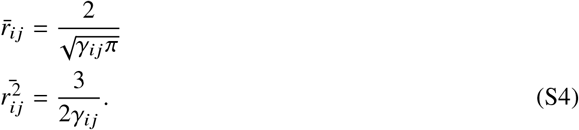

### 2 CONTACT PROBABILITY

Based on the above chromatin polymer model, we have discussed generic *n*-body contact probability in Ref. (21). The pairwise (*n* = 2) contact probability is given by

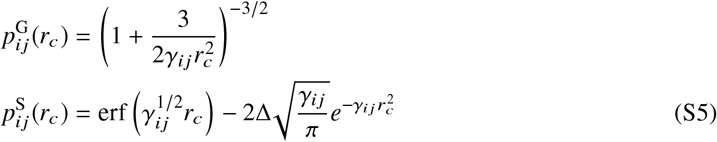

with the special function 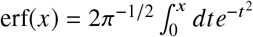. The superscript denotes the Gaussian or rectangular profile of *F*_Hi-C_(*r*), which describes the distance-dependent efficiency of the cross-linking agent in Hi-C experiments. More specifically, we have assumed

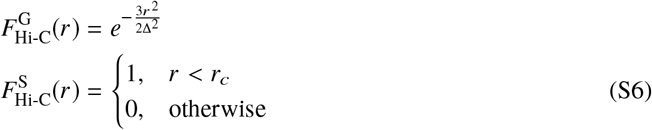

to obtain Eqs. S5 where Δ denotes a characteristic capture radius.

### 3 CO-SESEGREGATION PROBABILITY

To derive the co-segregation probability of chromatin segment pairs as the GAM experiment (14), we begin with the probability that the *i*-th segment is located at *z*_*i*_ = *α* from the COM of the chain,

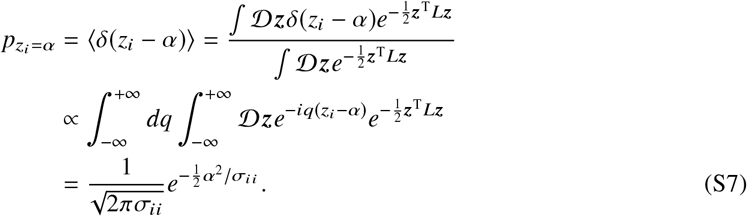

Similarly, the probability that the *i*-th and *j*-th segments are simultaneously found at *z*_*i*_ = α and *z*_*j*_ = *β*, respectively, is given by

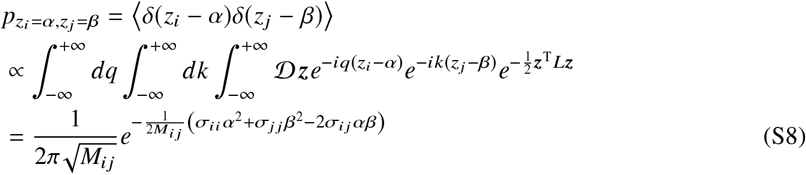

with 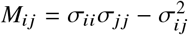.

#### Gaussian sectioning

Next, we assume that the horizontal slice 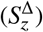 at *h* = *z* with thickness Δ is sectioned with a Gaussian probability

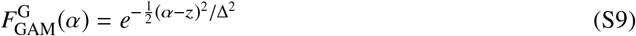

where the superscript again notates the Gaussian profile of *F*_GAM_. For simplicity, we have omitted the superscript “G” in the main text.

Combining Eqs. S7-S9, it is straightforward to formulate the probability of the *i*-th segment being sectioned by 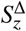 as

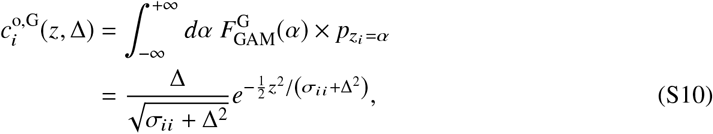

and the co-segregation probability as

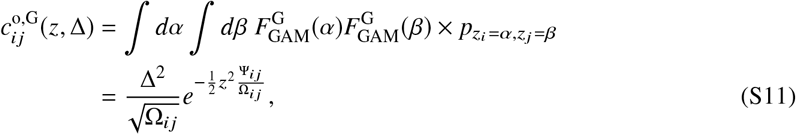

where

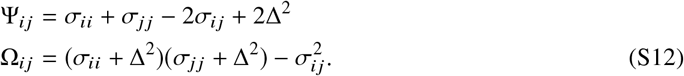

- **NPs collected from uniform slicing**. To mimic the procedure of GAM (14), we assume that the NPs are collected uniformly from *z* ∈ [−*R, R*] (*R* ≥ 0). Then the segregation probability is given by

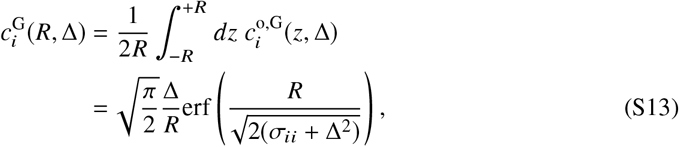

and the co-segregation probability has the expression of

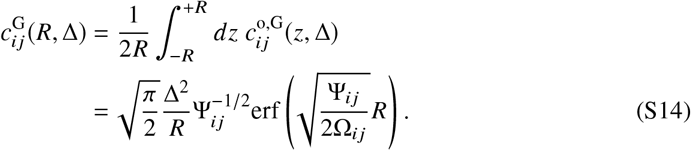
- **NPs collected from Gaussian slicing**. Alternatively, we assume the NPs are obtained from a Gaussian probability of 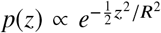 (i.e., a higher chance to be sliced at the center of cell nucleus than the apexes). This gives rise to the segregation probability as

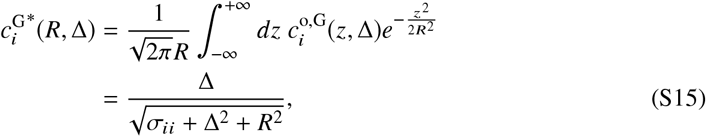

and the co-segregation probability as

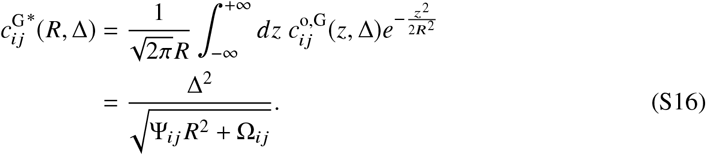

as shown in Fig. S2, 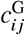 and its correlation with 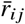 calculated using Eq. S16 are similar to those calculated with Eq. S14 (see Fig. 2D and E in the main text).

#### Rectangular sectioning

Sections with sharp edges in GAM can be modeled more realistically by employing a rectangular profile. The above results (Eqs. S10-S14) can be reformulated by replacing 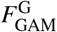 in Eq. S9 with

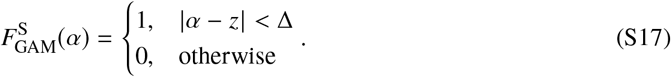

The segregation probability of the *i*-th segment from a slice made at *h* = *z* has a form of

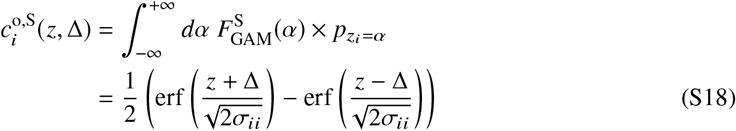

When the slices are collected from the range |*z*| ≤ *R*, the segregation probability becomes

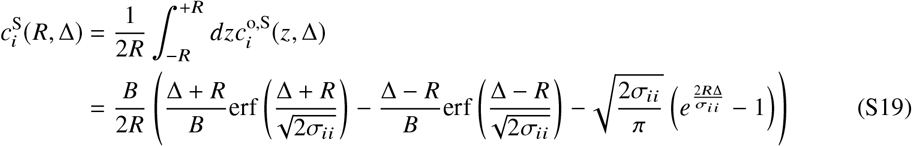

where 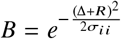. The corresponding co-segregation probability,

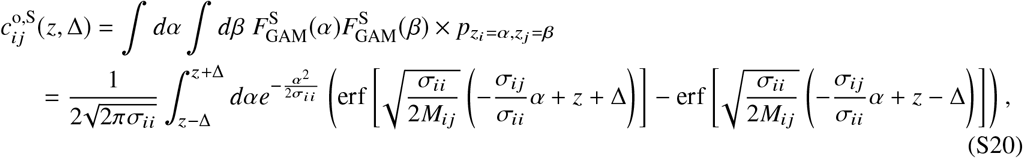

and 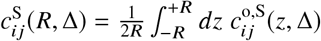 shown in Fig. S3 are obtained with numerical integration. All the results using rectangular sectioning are similar to those calculated using the Gaussian sectioning and are presented in Fig. S3.

### 4 3D STRUCTURES OF POLYMER CHAIN

In HLM, the normal coordinate vector of a polymer chain **X**_*α*_ = *Q*^T^***r***_*α*_ satisfies

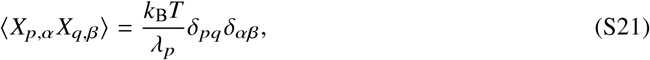

where *Q* and *λ* are defined in Eq. S2, *k*_B_*T* is our energy unit, *α* and *β* represent *x, y* and *z*, and *p, q* = 1, 2, 3, …, *N* − 1. Based on this relation, a 3D conformation of the polymer chain can be generated in two steps (43). First, we obtain the normal coordinates by using

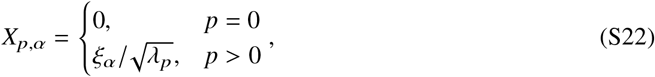

where the random variable *ξ*_*α*_ obeys the normal distribution with ⟨*ξ*_α_⟩ = 0 and 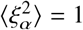. Next, the normal coordinates are converted to the Cartesian coordinates of polymer segments by

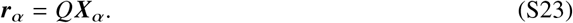

Repeating this protocol *n* times yields a structural ensemble of the polymer chain containing *n* samples.

**Figure S1:**
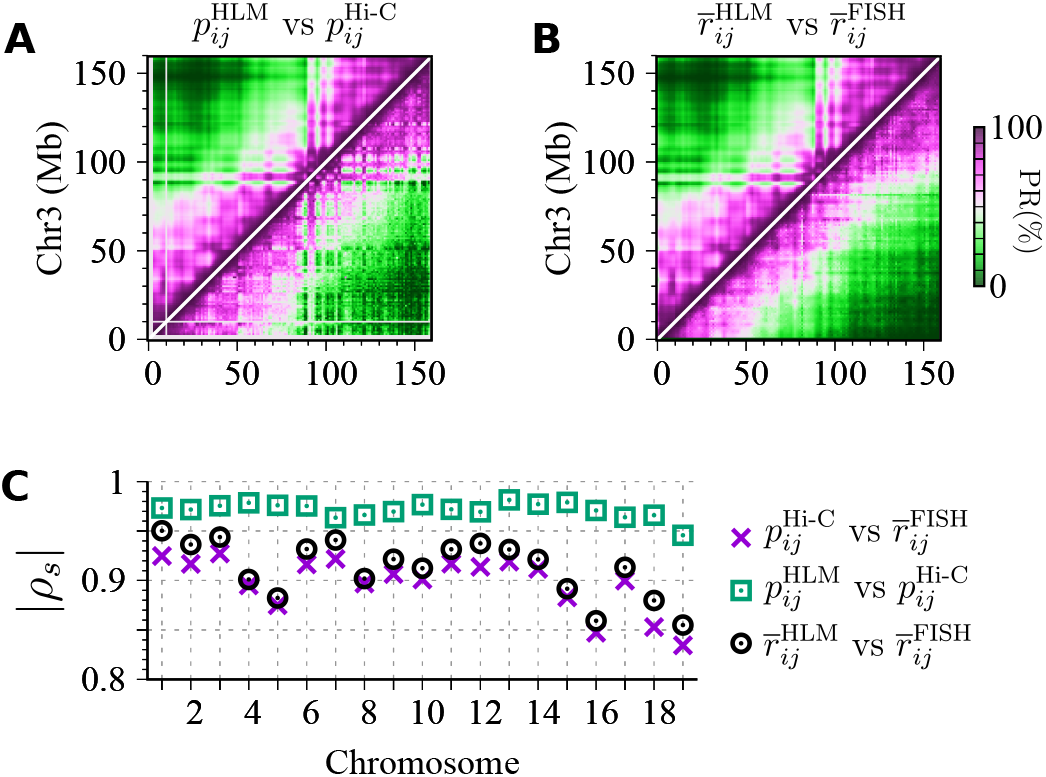
Comparing mouse ESC autosomes modeled at 1-Mb resolution with HLM against experimental data. (A) Percentile rank (PR) of the contact probability of chromosome 3 from HLM (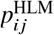, top left conner) versus that from Hi-C (5) (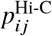, bottom right conner). (B) PR of the mean inter-locus spatial distance from HLM (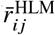, top left conner) versus that from FISH (24) (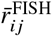, bottom right conner). (C) The Spearman’s rank correlation coefficient for all autosomes.

**Figure S2:**
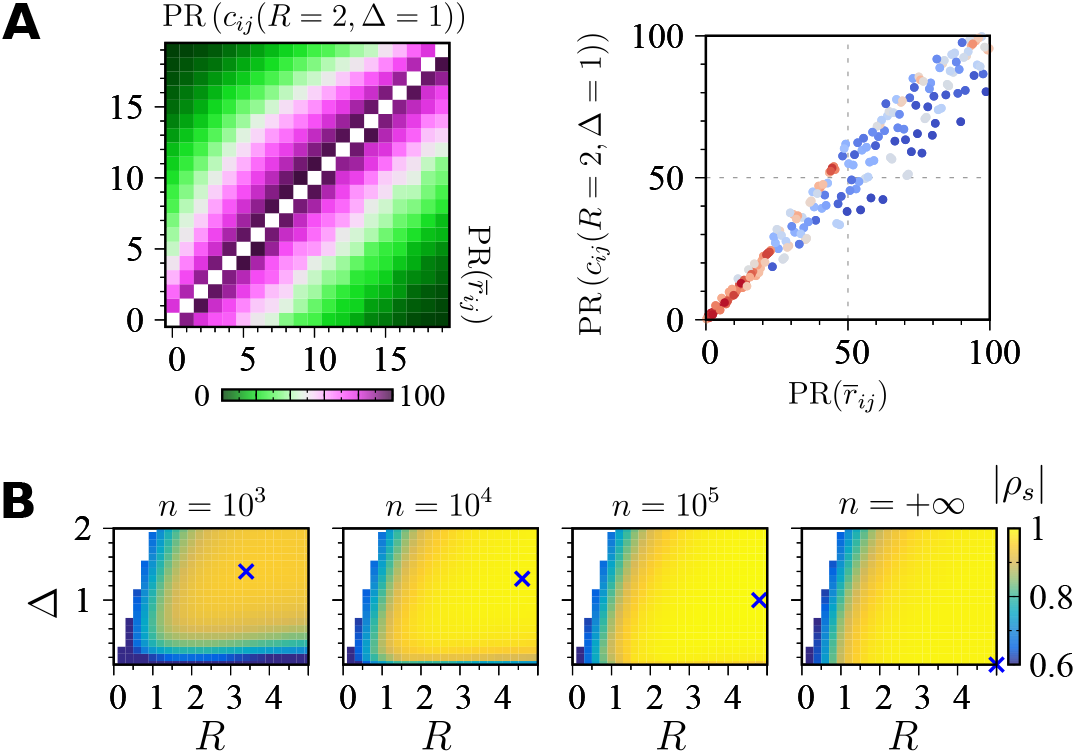
Co-segregation probability of a Gaussian polymer chain which are calculated by assuming the slices position following a Gaussian profile (see Eqs. S15 and S16). **(A)** and **(B)** are similar to Fig. 2D-E in the main text, respectively.

**Figure S3:**
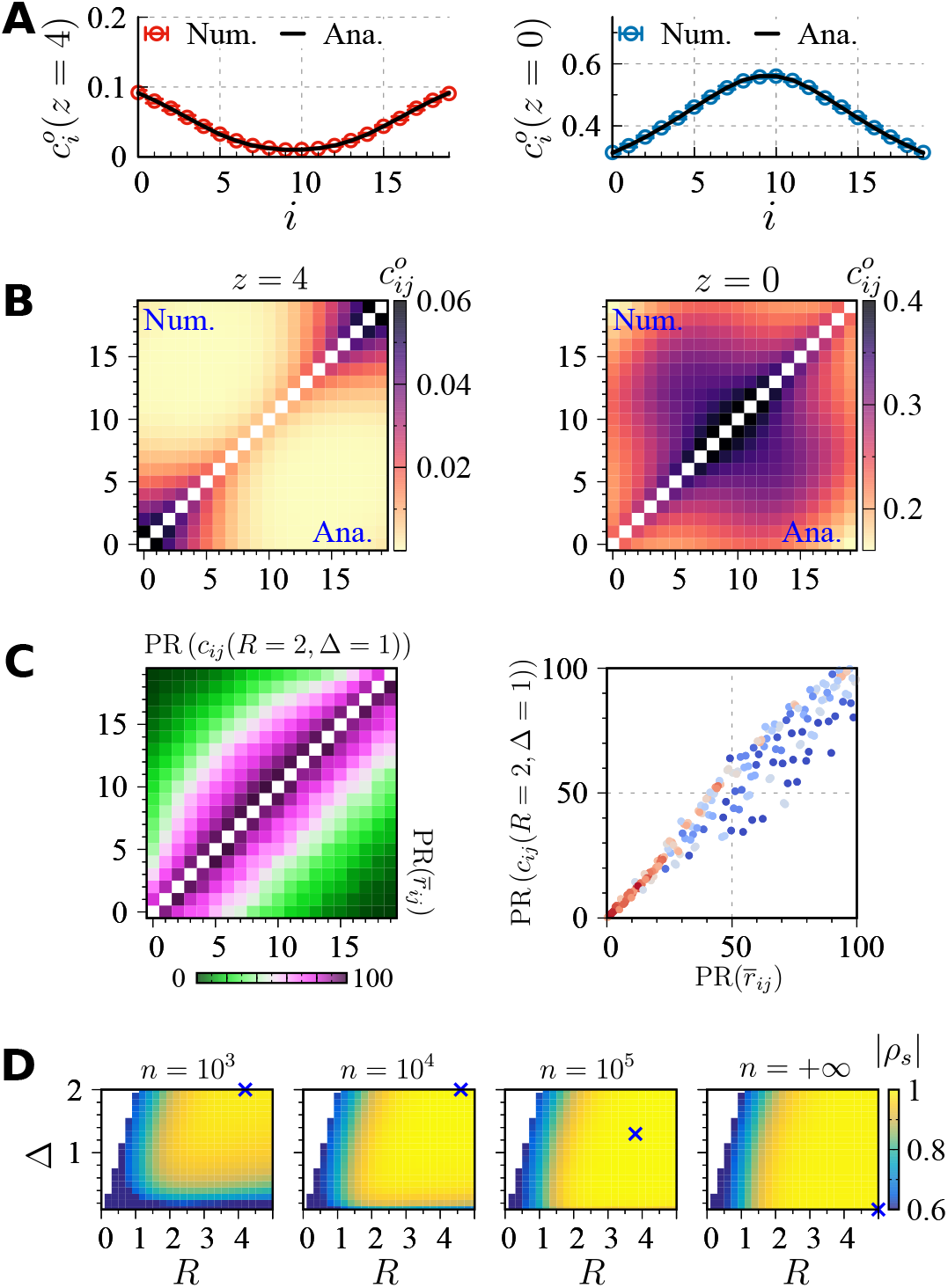
Co-segregation probability of a Gaussian polymer chain which are calculated by assuming the rectangular sectioning profile (see Eqs. S17-S20). **(A)**-**(D)** are similar to Fig. 2B-E in the main text, respectively.

**Figure S4:**
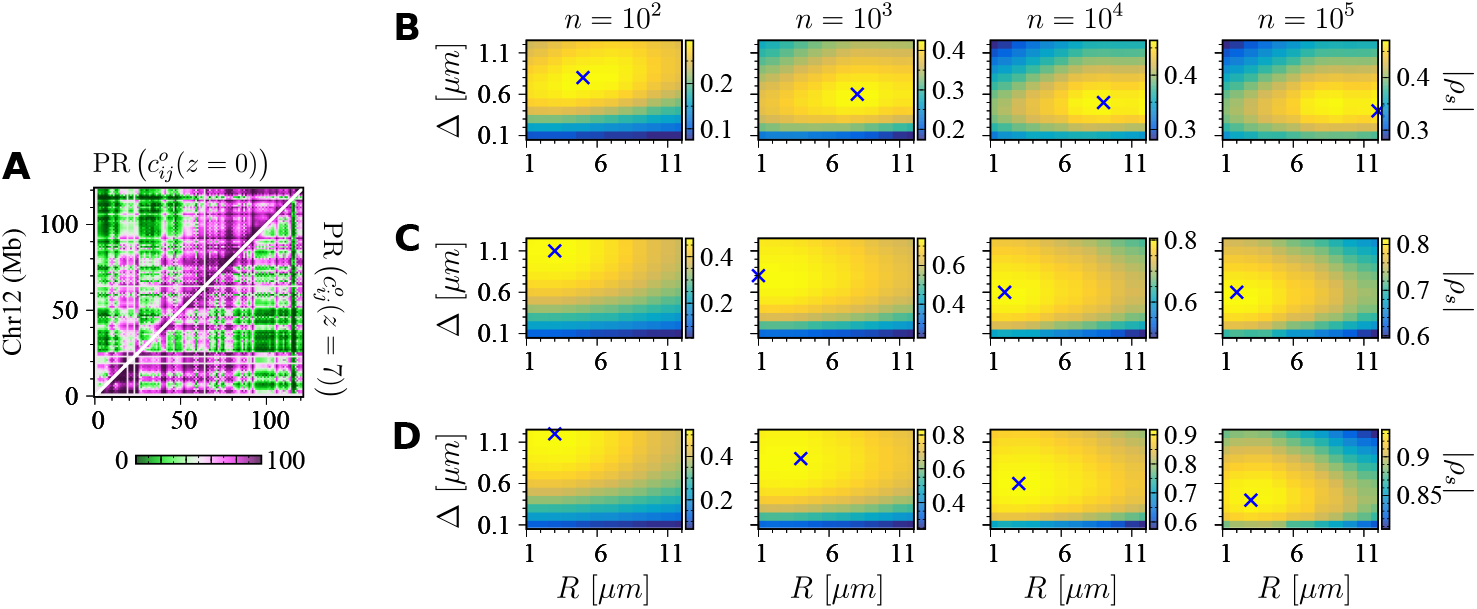
In silico GAM results of chromosome 12. **(A)** Percentile ranks of *c*_*ij*_ sectioned by slices with a width of Δ = 0.5 *µ*m and a height of *z* = 0 (top-left) and 7 *µ*m (bottom-right). The Spearman’s correlation coefficient between the two is *ρ*_*s*_ = 0.75. **(B)** The correlations (ρ_*s*_) between the mean physical distance and co-segregation probability, **(C)** NLD, and **(D)** NPMI as a function of the positional uncertainty (*R*) and width (Δ) of slices for varying *n*. The symbol “×” denotes the values of (*R*, Δ) that maximizes |*ρ*_*s*_ |.

**Figure S5:**
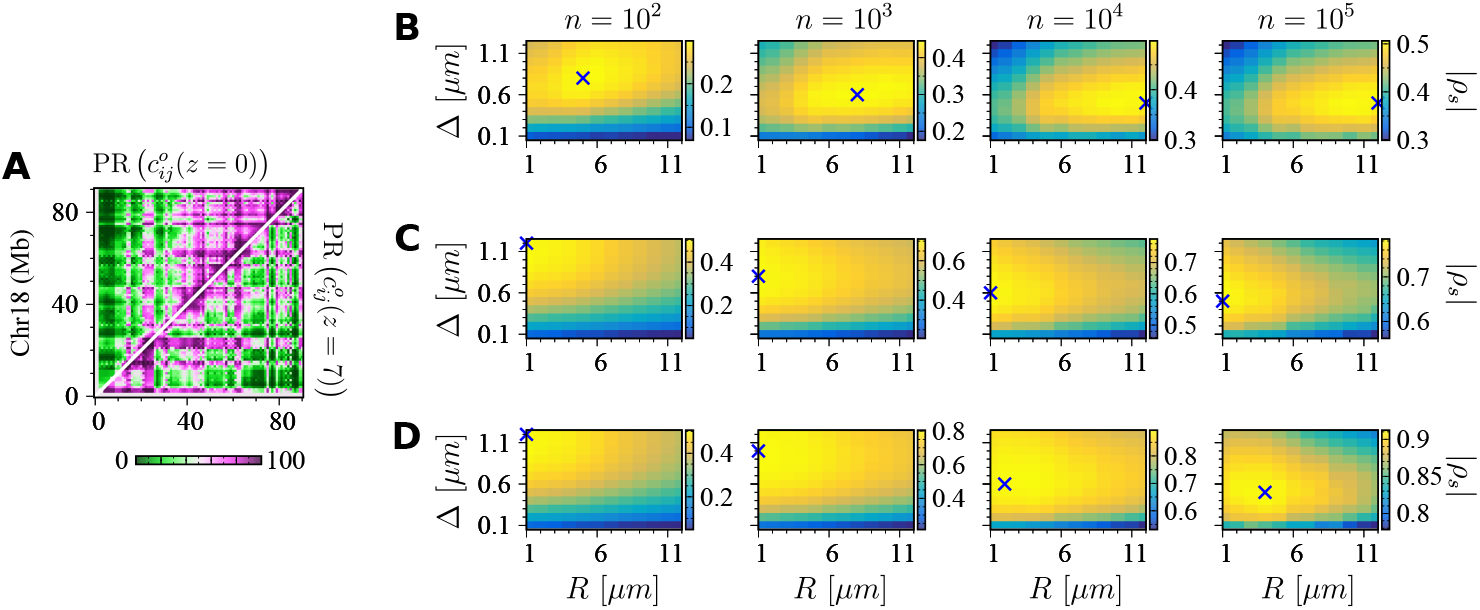
In silico GAM results of chromosome 18. Please refer to the caption of Fig. S4 for explanations. The Spearman’s correlation between the top-left and bottom-right panels in **(A)** is 0.69.

**Figure S6:**
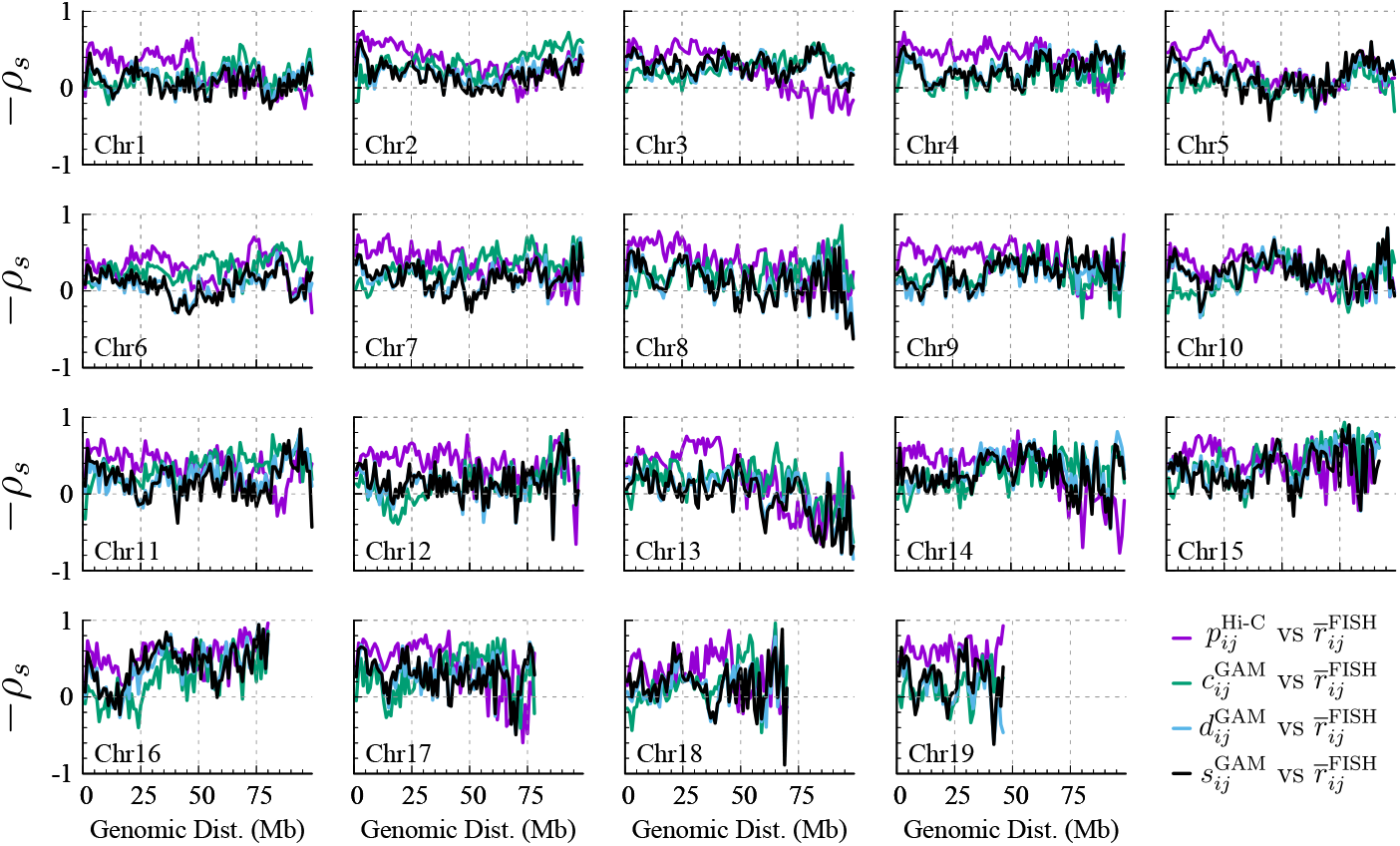
The Spearman’s correlation between the mean physical distance 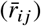 and Hi-C contact probability (*p*_*ij*_), GAM co-segregation probability (*c*_*ij*_), and its two variants (*d*_*ij*_ and *s*_*ij*_) as a function of the genomic distance between the (*i, j*) sites in different chromosomes. The mean and standard deviation averaged over all the autosomes are shown in Fig. 5A in the main text.

